# Helix: a structure-aware deep learning model for accurate prediction of A-to-I RNA editing by endogenous ADARs

**DOI:** 10.64898/2025.12.18.695251

**Authors:** Yingxin Cao, Lina R. Bagepalli, Yiannis A. Savva, Sierra M. Collins, Kristofor Adams, Alexander J. Letizia, Joanne Boysen, Aydin Abiar, Samantha R. Edwards, Lillian Bussema, Stephen M. Burleigh, Lucia Shumaker, Ronald J. Hause, Brian J. Booth, Yue Jiang

**Author notes:** Corresponding authors: Brian Booth and Yue Jiang.

## Abstract

Adenosine deaminase acting on RNA (ADAR) converts adenosine to inosine within double-stranded RNA (dsRNA) and can be co-opted for therapeutic RNA editing by introducing dsRNA substrates in trans using programmable guide RNAs (gRNAs). However, ADAR’s natural promiscuity necessitates sophisticated gRNA designs to achieve efficient and specific editing. Here we present Helix, a predictive model that achieves highly accurate, zero-shot per-adenosine editing predictions for any target sequence. Helix’s performance arises from two architectural choices: a transformer framework that scales effectively with increasing and imbalanced training data; and a structure-aware attention mechanism that incorporates predicted RNA secondary structure, a key determinant of ADAR activity. Helix’s predictive accuracy enables seamless integration with DeepREAD, our previously reported generative model, in a noisy-student distillation framework termed DeepHelix. This workflow supports both zero-shot gRNA design and the generation of complex, constraint-based designs. We demonstrate DeepHelix’s utility by designing gRNAs that efficiently edit a therapeutically relevant adenosine and by leveraging its flexible design space to engineer species cross-reactive gRNAs to accelerate pre-clinical development.

---

**A**denosine deaminase acting on RNA (ADAR) is a ubiquitously expressed enzyme that converts adenosine to inosine within double-stranded RNA (dsRNA), where inosine is interpreted as guanosine during splicing and translation^1–3^. By delivering guide RNAs (gRNAs) that hybridize to form a dsRNA structure with a target transcript, ADAR can be recruited to modulate regulatory motifs, recode amino acids to alter protein function, or correct pathogenic G-to-A mutations in a programmable manner^4–9^. A major challenge for therapeutic RNA editing is ADAR’s natural promiscuity and the impact of local sequence and secondary structure^10,11^. Chemically synthesized gRNAs can incorporate base, sugar, and backbone modifications to improve editing efficiency and specificity^12,13^, but DNA-encoded gRNAs that allow one-time therapeutic administration, such as those delivered by adeno-associated virus (AAV), must rely entirely on the secondary structure formed upon hybridization of the gRNA and target RNA^14–16^. As a result, accurate prediction of ADAR editing outcomes requires models that are sensitive to both sequence context and RNA secondary structure.

To address this challenge, we previously reported DeepREAD^17^, a generative bit-diffusion framework combining high-throughput screening (HTS) and machine learning to design gRNAs for unseen targets. DeepREAD performed well when trained on a balanced library containing thousands of target RNAs, but did not improve when trained on additional, highly imbalanced datasets, such as large single-target libraries containing > 10^3^ gRNAs. In contrast, large language models (LLMs) are expected to continually improve with additional data^18,19^. This reflects a broader trend in biological modeling: while LLMs often benefit from scaling laws^20–24^, other applications frequently require carefully curated data and task-specific architectures to achieve optimal performance^25–28^.

In drug discovery, experimental data are frequently imbalanced and target specific. We therefore sought a model that (i) improves with additional HTS datasets even if they are imbalanced, (ii) explicitly incorporates the sequence and structural determinants of ADAR editing, and (iii) benefits from fine-tuning on small datasets that are often generated as gRNA candidates advance into disease-relevant cellular systems. In parallel, we aimed to integrate prediction with generative design in a single iterative workflow.

To meet these needs we developed Helix, which has a custom transformer-based architecture^29^ and a modified attention mechanism that incorporates predicted RNA secondary structure. Helix scales effectively with large, imbalanced screening datasets and achieves high performance on ADAR editing prediction. We then paired Helix with DeepREAD in a noisy student (NS) distillation framework^30^ to create DeepHelix: a unified predictive–generative workflow that enables knowledge transfer between models, structurally informed gRNA designs, and post-generation filtering to prioritize gRNAs with the desired attributes. The workflow combines the strengths of both models through three steps: (i) using Helix to score candidate gRNAs and provide functional pseudo-labels for DeepREAD training; (ii) using DeepREAD to generate large pools of candidate gRNAs; and (iii) using Helix again to score, rank, and filter these candidates for in-cell testing. This workflow supports both zero-shot inference and more complex design tasks, such as species cross-reactivity for preclinical development. DeepHelix provides a flexible, biologically informed platform for rapid gRNA design and iterative optimization tailored to therapeutic RNA editing.

## Results

### Helix architecture enables efficient learning from imbalanced RNA editing datasets

We developed Helix, a transformer architecture designed to predict ADAR-mediated RNA editing outcome at per-adenosine resolution while efficiently leveraging imbalanced training datasets (Fig. 1). Helix uses an encoder–decoder transformer with 20 attention heads. One-hot encoded gRNA–target sequences with positional encoding pass through the encoder, and the resulting representations are split into gRNA and target components before entering the decoder, which produces position-specific editing predictions along the gRNA hybridization region of the target sequence, including the on-target and bystander adenosines. Summary statistics including on-target editing efficiency, specificity (log ratio of on-target over total bystander edits) and maximum bystander editing can then be derived from per-adenosine editing predictions.

**Fig. 1:**
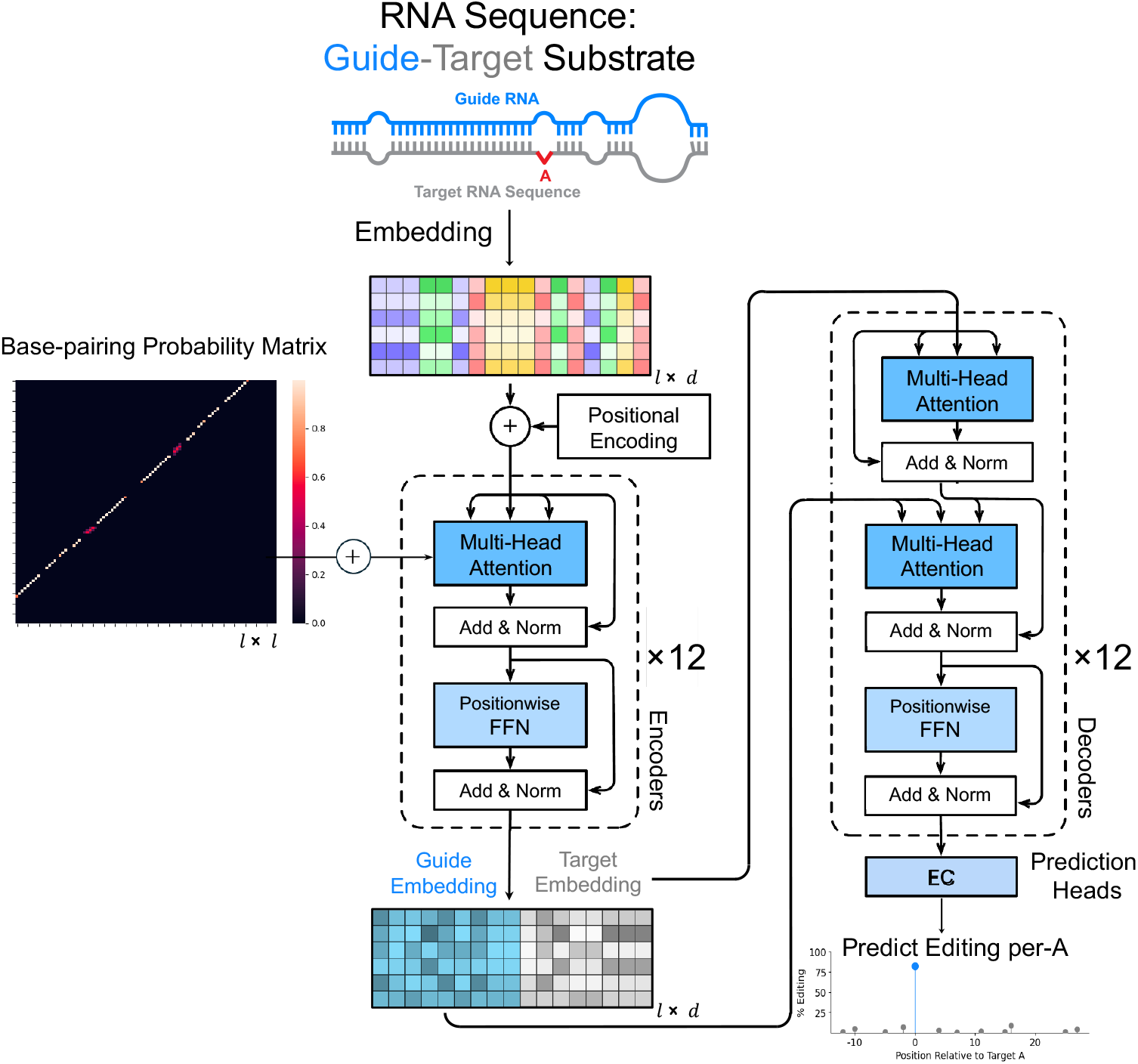
Helix Model Architecture. Helix is a transformer-based sequence-to-sequence deep learning model for predicting outcomes of ADAR-mediated RNA editing. The model takes one-hot encoded RNA sequences of concatenated gRNA-target pairs as input, and outputs a numerical prediction track of editing efficiencies for each adenosine within the gRNA hybridization region of the target sequence. The encoder block consists of 12 transformer encoder layers with modified self-attentions. In Helix, the standard self-attentions are augmented by additions of Base-pairing Probability Matrices (BPMs) at each layer to explicitly represent the secondary structures of the target-gRNA duplex. The resulting intermediate embeddings are then split into target and guide components and passed to the decoder, where the target embedding serves as the query (Q) and the guide embedding serves as keys and values (K, V) in 12 layers of cross-attention. A final linear prediction head outputs editing probabilities at single-nucleotide resolution.

Our first objective was to determine how transformer models respond to imbalanced training data. The available datasets span three levels of target-to-gRNA sampling density (Fig. 2a): the PolyTarget library samples a total of 5,643 targets with an average of only 10 gRNAs per target; the DeepREAD validation libraries contain 42 targets with up to 2,000 gRNAs per target; and five single-target HTS libraries with approximately 20,000 gRNAs per target. Together, these datasets demonstrate the tradeoff between target and gRNA sampling densities.

**Fig. 2:**
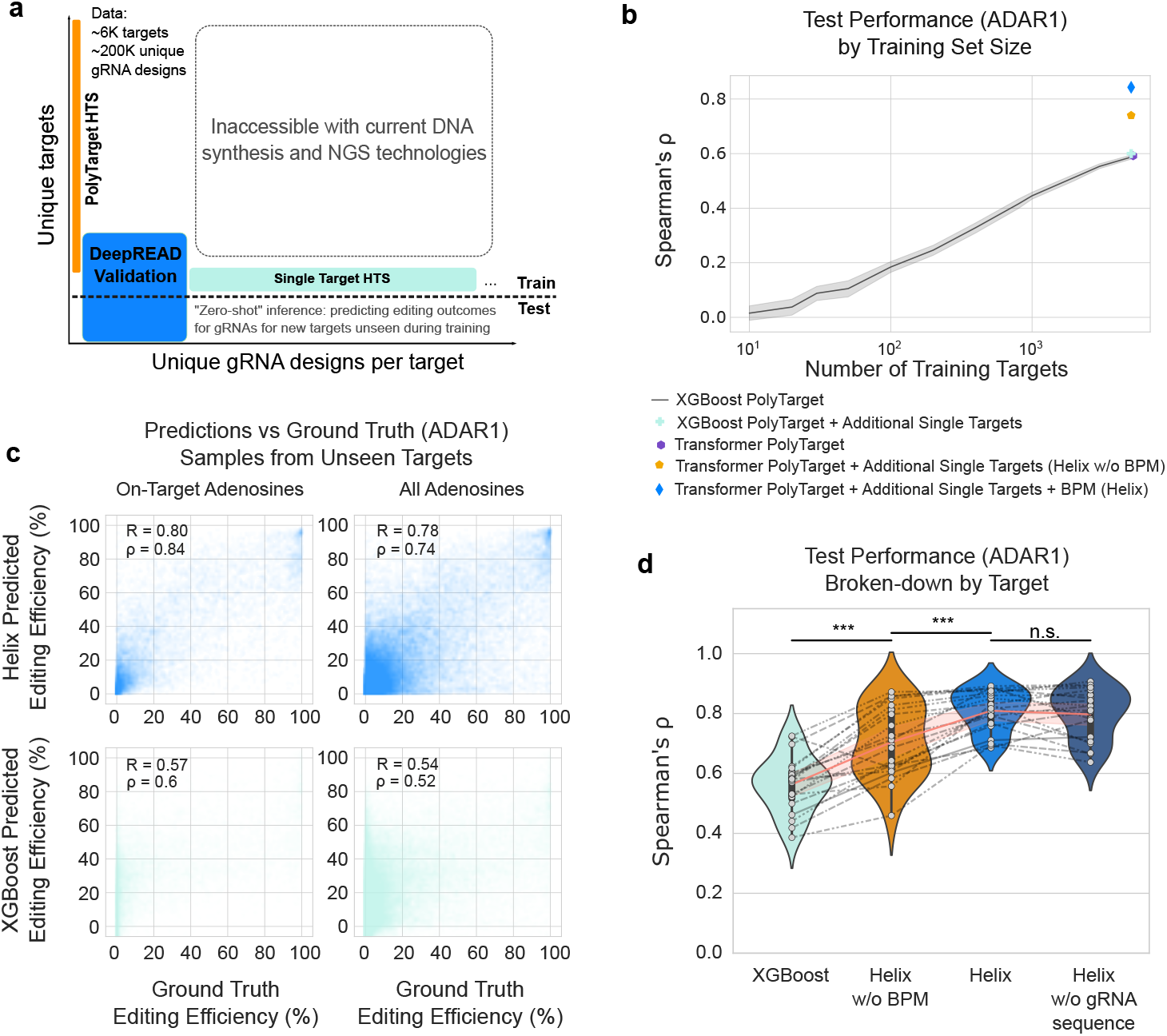
Datasets and model performance. **a**, Composition of training and testing data. Training data contains samples from ∼6,000 therapeutically relevant targets, with ∼200,000 unique gRNAs in total. It contains all the data from PolyTarget library and single-target HTS libraries, and part of the DeepREAD validation libraries where targets overlap with the PolyTarget or single-target libraries. The test data consists of ∼20,000 gRNA designs for 21 unseen targets from the DeepREAD validation library. **b**, Spearman correlation between test predictions and ground truth for ADAR1 editing outcomes across models, as a function of the number of targets used for training. **c**, Predictive performances on ADAR1 editing by Helix and XGBoost for on-target positions, and all adenosine containing positions (bystander and on-target). **d**, Spearman correlation between predictions and ground truth for ADAR1 editing at on-target positions averaged within each target across different models. ^***^ indicates p value < 0.01.

To evaluate how model performance scales with additional training data, we withheld 21 targets (36,000 gRNA–target pairs) from the DeepREAD validation library as a zero-shot test set and progressively expanded the training data from PolyTarget to include the validation and single-target libraries. Predictive performances were measured by Spearman correlation (ρ) between predicted and experimental on-target editing (Fig. 2b, Supplementary Fig. 1a). As more targets from the PolyTarget library were added, both XGBoost and transformer models improved to ρ ≈ 0.6, consistent with our previous data^17^. Interestingly, when high-density single-target libraries were incorporated, the transformers continued to improve, reaching ρ = 0.74, while XGBoost showed no additional gains. This indicates that transformer models are uniquely capable of leveraging densely sampled, structurally diverse gRNA data from a small number of targets and generalizing to unseen sequences.

### Integration of secondary structure information significantly enhances model performance

RNA secondary structure contains important information predictive of ADAR editing outcomes^31^, and current structure prediction tools achieve high accuracy^32,33^. To assess if the transformer model learns such structural information, we examined attention maps from the final encoder layer, which reflect the model’s internal representation of the gRNA–target duplex. Regions of high attention scores aligned with structured portions of the duplex, indicating the model infers relationships between primary sequence and predicted secondary structure. However, high attention scores are also found near the flexible AAAAA linker inserted between target and gRNA to facilitate in silico folding – an artificial feature present in all designs that provides little biological information beyond marking the junction between the two sequences (Fig. 3d). These observations led us to hypothesize that providing the model with explicit secondary-structure information within the attention mechanism could further improve predictive performance.

**Fig. 3:**
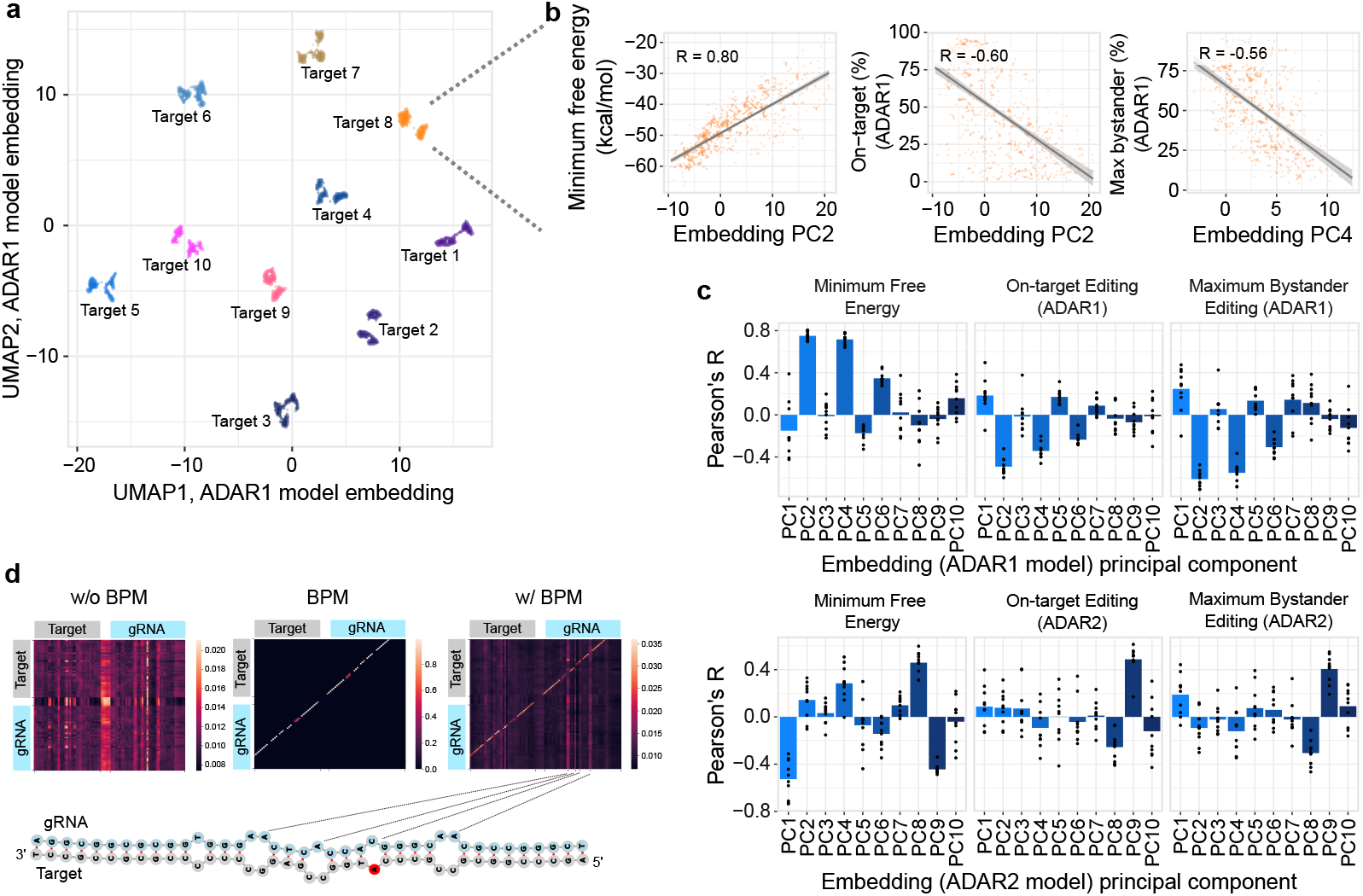
Helix captures important target and gRNA features related to editing. **a**, 2-dimensional UMAP visualization of embeddings from the transformer encoder block of Helix ADAR1 model, for 10 targets with 600 gRNAs each. Each dot represents a gRNA sample. **b**, Examples of top principal components (PC) of embeddings in relation with target-gRNA substrate features including minimum free energy (MFE), on-target editing and maximum bystander editing. **c**, Pearson correlation coefficients of the top 10 PCs of ADAR1 and ADAR2 embeddings with MFE, on-target editing and maximum bystander editing. **d**, Heatmap of attention scores from the last layer of the transformer encoder produced by Helix without and with the addition of the base-pairing probability matrix, for an example target-gRNA duplex. The predicted structure of the sequence is shown below, with arrows indicating positions with secondary structural features.

To test this hypothesis, we predicted the secondary structure of each target-gRNA duplex^33^ and supplied the resulting base-pairing probability matrix (BPM) as a numerical mask for attention (see Methods). Adding the BPM biases the attention toward nucleotide pairs with high predicted pairing probability, thereby exposing the model directly to structural information. Explicitly providing this structural prior led to a marked improvement in predictive performance (Fig. 2b, Spearman’s ρ = 0.84, Supplementary Fig. 1a). When evaluated across all adenosines on the target, the model also achieved high accuracy (Fig. 2c, Spearman’s ρ = 0.74, Supplementary Fig. 1b). Both metrics significantly exceed XGBoost models trained and evaluated using the same data (Spearman’s ρ = 0.60 for on-target editing and 0.52 for all adenosines).

To further evaluate the role of secondary structure, we performed an ablation study in which the gRNA sequence was removed from the input, leaving the model solely trained on target sequence and predicted secondary structure of the target-gRNA duplex. For many targets the gRNA ablated model performed comparable to the full model, indicative of the key role that secondary structure plays in ADAR editing (Fig. 2d, Supplementary Fig. 1c). Hereafter, Helix refers to the transformer model trained on all datasets with primary sequences and secondary structures. Separate Helix models were trained for ADAR1 and ADAR2 editing.

### Helix captures important target and gRNA features

Transformers represent each nucleotide position as a continuous embedding vector that integrates sequence identity, positional context, and information aggregated through attention. Given that the BPM biases the attention mechanism toward paired nucleotides, the embedding space is a natural place to examine whether Helix has internalized secondary-structure cues. To do so, we calculated the target-gRNA embeddings for 600 gRNAs per 10 targets using Helix ADAR1 and ADAR2 models. The embeddings clearly separate by target when visualized using two-dimensional Uniform Manifold Approximation and Projection (2D UMAP) (Fig. 3a, Supplementary Fig. 2a). Focusing on single targets, the principal components (PCs) of embeddings show various degrees of correlation with minimum free energy (MFE), a univariate continuous surrogate for secondary structures of the target-gRNA duplex. Importantly, one can also find PCs that are strongly correlated with predicted editing outcomes, including on-target editing and maximum bystander editing (Fig. 3b, c, Supplementary Fig. 2b). These observations suggest that embeddings obtained from the encoder layers of Helix contain important information about the secondary structures and ADAR editing outcomes, as opposed to a black box that is uninterpretable. For example, PC2 has a strong positive correlation with MFE and a strong negative correlation with on-target editing (Fig. 3b). This inverse trend reflects ADAR’s role as a dsRNA binding and editing enzyme. Increasing the number of unpaired bases (i.e., secondary structures) will increase the MFE and reduce ADAR binding and activity. The same trend exists for bystander editing, highlighting the balance required to utilize secondary structures to minimize bystander editing while maintaining on-target editing. The strong correlation between simple linear transformations of embedding as represented by PCs and predicted editing summary statistics also confirms that more complex linear transformations of embeddings from Helix as performed by the decoders are expected to provide accurate predictions for editing outcomes. Overall, these observations suggest that the embeddings from Helix have learned information about RNA secondary structures and ADAR activity.

### Knowledge distillation from Helix enhances generative gRNA design

We previously reported DeepREAD, a generative bit-diffusion model that produces highly efficient and specific gRNAs for unseen targets^17^. Although DeepREAD performs well on the balanced PolyTarget HTS library, it failed to scale when large single-target HTS datasets were added. This is likely because the extreme class imbalance in single-target libraries induced mode collapse, a well-documented challenge for generative models when rare classes become poorly represented^34,35^. Given this limitation, we propose a new workflow, DeepHelix, for generative gRNA design for unseen targets leveraging NS distillation to pair the predictive power of Helix alongside DeepREAD’s generative design framework (Fig. 4a). In this setup, tens of thousands of unlabeled gRNAs to one or more targets are randomly generated by programmatically introducing various secondary structural features at user defined distributions to the seed sequence, which by default is the reverse complement of the target sequence (see Methods). Helix is then used as the teacher model to score all these gRNA designs to form a large dataset with pseudo-labels. We then add noise to the Helix predictions and use it to train a NS bit-diffusion model. The diffusion model trained on pseudo-labels is used to generate gRNA design candidates conditioned on desired editing properties. The generated diffusion samples were then scored by Helix again, pooled alongside the initial random in silico designs, to go through a final round of prediction and filtering to prioritize gRNA candidates for in-cell testing. As expected, the DeepHelix workflow was able to generate more gRNA designs with Helix-predicted high efficiency and specificity compared to the initial random designs (Fig. 4b).

**Fig. 4:**
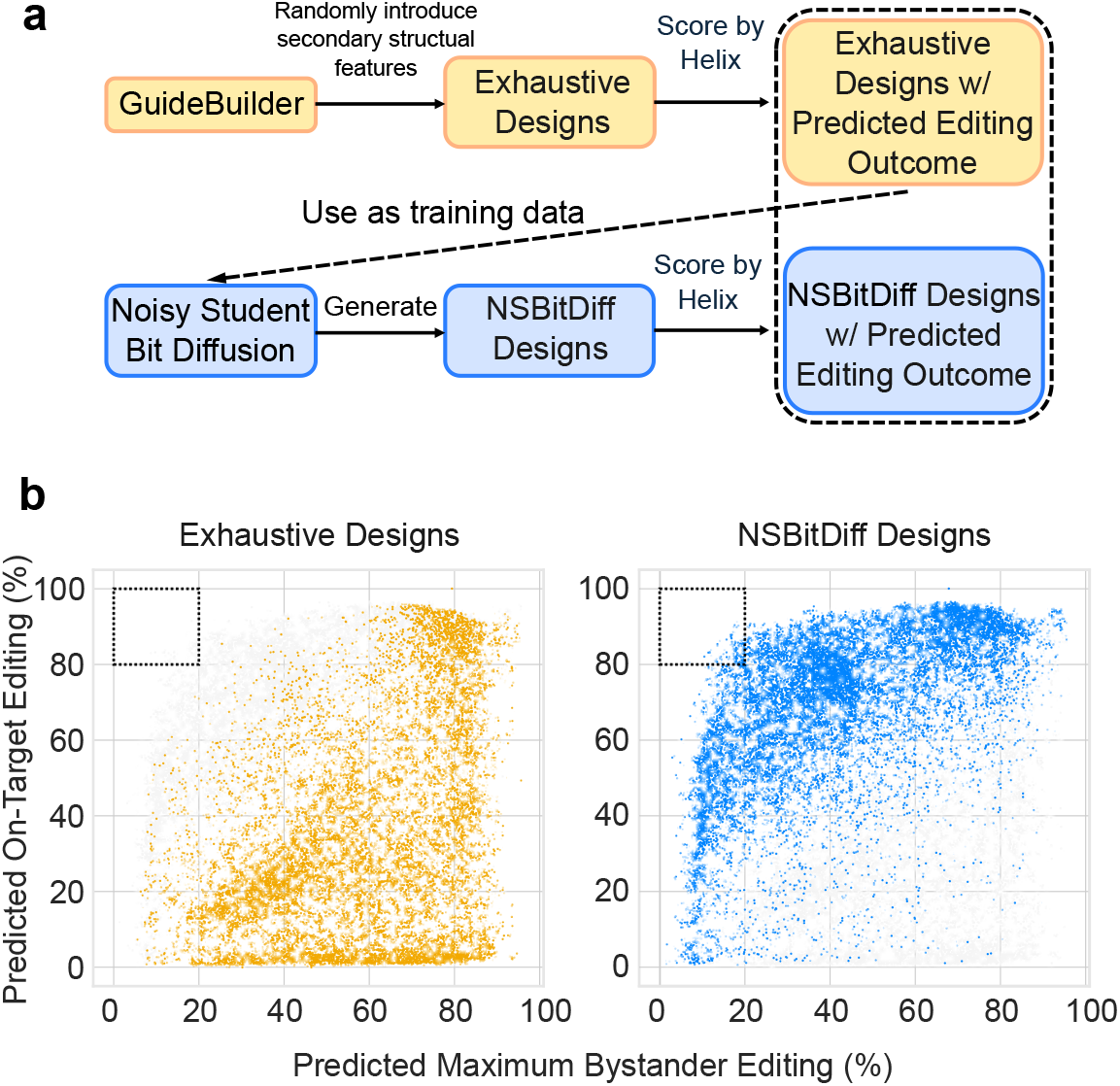
Knowledge distillation with noisy student for gRNA design. **a**, DeepHelix workflow for noisy student (NS) distillation. Tens of thousands of random gRNAs are generated using GuideBuilder by introducing secondary-structure features at user-defined distributions into a seed sequence. Helix predicts editing outcomes for these designs, and noise is added to these predictions to create pseudo-labels for training the NS bit-diffusion model. The diffusion model then generates new gRNA candidates conditioned on desired editing properties. These diffusion-generated samples are rescored by Helix and pooled with the initial random designs for a final filtering step that prioritizes tens to hundreds of candidates for in-cell validation. **b**, Helix-predicted editing efficiencies and maximum bystander editing for exhaustive random designs (left) and NS bit-diffusion designs after NS distillation (right).

### Zero-shot design and screening of therapeutic gRNAs in cells

DeepHelix was used to generate gRNAs targeting the adenosine within the translation initiation site (TIS) of the GRIK2 transcript. A HEK293T GRIK2 minigene model (293-hGRIK2) confirmed that converting the TIS **A**TG-to-**G**TG abolishes translation of the encoded GluK2 protein (Supplementary Fig. 3). Reducing GluK2 levels may provide therapeutic benefit for patients with mesial temporal lobe epilepsy (MTLE), a disease associated with aberrant GluK2 expression^36^.

Previous studies have shown that DNA-encoded gRNAs must exceed ∼80 nucleotides to efficiently recruit endogenous ADAR^8,9^. However, long hairpin substrates present practical challenges for DNA synthesis, reverse transcription, and PCR. As a result, our training data were generated from a biochemical assay using in vitro–transcribed RNA hairpins of ∼45 base pairs. The shorter gRNAs produced by DeepHelix were therefore extended to ∼100 nucleotides for in-cell validation (see Methods).

We screened 100 DeepHelix-designed gRNAs in 293-hGRIK2 cells and identified 12 candidates with ≥50% on-target editing and ≤15% maximum bystander editing (Fig. 5a). These high performing gRNAs displayed diverse sequence composition and secondary structures (Supplementary Fig. 4), which helps de-risk sequence dependent liabilities that may arise during pre-clinical development^37^. The per-adenosine editing profiles predicted by Helix closely matched experimental measurements (Supplementary Fig. 4b), demonstrating both the accuracy and the fine-grained resolution of Helix.

**Fig. 5:**
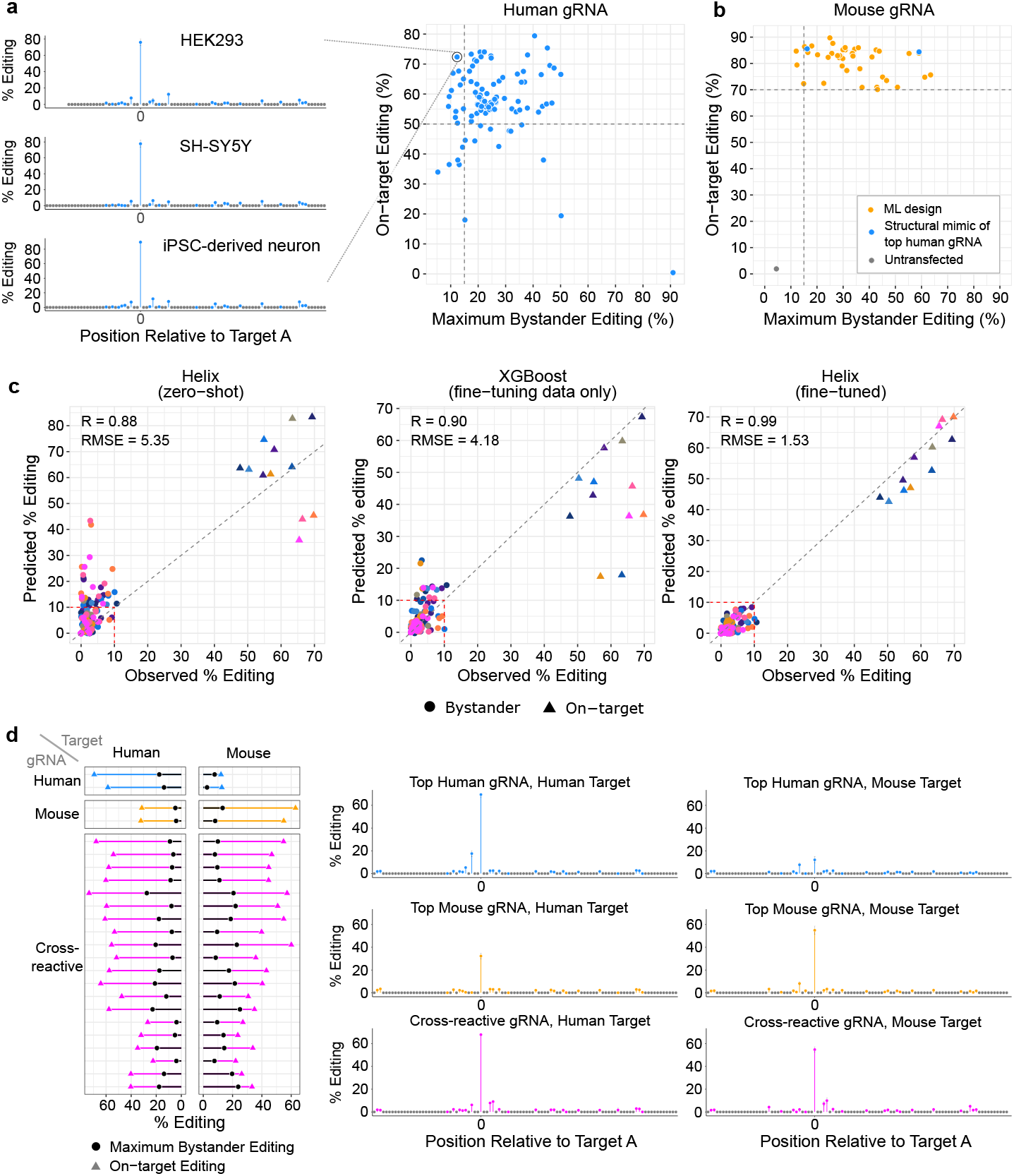
In-cell validation of Helix designs and extended use cases using GRIK2 TIS as an example. **a**, Validation of gRNAs designed for GRIK2 TIS in the noisy student distillation procedure in HEK293 cells. Example editing profiles of a top gRNA candidate with high on-target editing efficiency and low bystander editing in HEK293 cells, SH-SY5Y cells and iPSC-derived neurons. **b**, Screening of surrogate gRNAs designed for mouse Grik2 TIS in HEK293 cells. **c**, Fine-tuning Helix enables further diversification of top candidate gRNAs. Predictions of editing performance of the second round of gRNA designs by Helix (zero-shot) without fine-tuning, XGBoost only trained on fine-tuning data, and fine-tuned Helix. **d**, Designing gRNAs with cross-species reactivity. Editing performance of two top human GRIK2 TIS gRNAs, two mouse Grik2 TIS gRNAs and 20 gRNAs designed for cross-species reactivity, assayed on both the human and mouse targets. Example editing profiles of one top candidate for the human target, mouse target and both targets are highlighted.

A subset of these candidates was packaged into the AAV-DJ serotype for broad cell-type tropism^38^ and delivered to induced pluripotent stem cells (iPSC) derived neurons and differentiated SH-SY5Y cells to assess editing of the endogenous GRIK2 transcript. These gRNAs achieved similarly efficient and specific editing across both neuronal systems, reaching > 90% on-target editing in iPSC-derived neurons after 10 days and > 75% in SH-SY5Y cells after 7 days (Fig. 5a). These results demonstrate that the DeepHelix workflow enables true zero-shot design of highly active gRNAs in biologically relevant cell types.

### DeepHelix identifies species cross-reactive gRNAs

In contrast to the result seen in human cells, the top human gRNAs showed poor cross-reactivity with the mouse Grik2 TIS sequence, reflecting the fundamental difficulty of forming ADAR-compatible duplex structures across divergent sequences using a single gRNA (Fig. 5d, Supplementary Fig. 5b). Homology across relevant animal models is a key consideration during pre-clinical development^39^. The human GRIK2 TIS target region shares 85% sequence identity with mouse Grik2 (Supplementary Fig. 5a), which presents a design challenge for ADAR substrates that are highly sensitive to small secondary-structure perturbations^16^. To evaluate whether DeepHelix could overcome this challenge, we first used zero-shot inference to design mouse-specific surrogate gRNAs targeting the mouse GRIK2 TIS. We screened 40 DeepHelix-generated candidates in HEK293T cells expressing a mouse Grik2 minigene and identified multiple gRNAs with ≥70% on-target editing, including three with ≤15% maximum bystander editing (Fig. 5b). Although some structural mimics of top human gRNAs functioned on the mouse substrate, the generative approach produced a broader and more diverse set of mouse-active candidates. However, none of the mouse-specific designs showed strong cross-reactivity with the human target.

The flexibility of the DeepHelix workflow makes it well suited for engineering species cross-reactive gRNAs. To promote hybridization to both human and mouse targets, we introduced mutations at divergent positions within the seed region that allow either canonical or wobble base-pairing with both sequences. We then introduced additional randomized structural features to generate a diverse set of candidate gRNAs, which Helix scored against both the human and mouse targets. These Helix-labeled sequences were used to construct NS training samples with increased predicted affinity for both species. Training the DeepREAD on this dual-species dataset skewed the model’s sequence distribution towards cross-reactive designs and enabled it to generate gRNAs with high predicted efficiency and specificity for both human and mouse GRIK2 TIS sequences. Twenty DeepHelix generated cross-reactive gRNAs were tested experimentally on both targets, and several exhibited editing efficiencies approaching those of the best human- or mouse-optimized gRNAs (Fig. 5d). These results highlight the flexibility of DeepHelix to incorporate custom constraints and condition the generative process on multi-species performance, enabling rational design in settings where species cross-reactivity is desired for pre-clinical development.

### Fining-tuning Helix with minimal in-cell data improves performance

Drug discovery often requires iterative design cycles in which initial experimental results guide subsequent optimization^40^. To evaluate whether Helix can incorporate small in-cell datasets for target-specific refinement, we fine-tuned the model using 62 in-cell editing measurements for gRNAs targeting the GRIK2 TIS. Many initial gRNA designs achieved high editing efficiency but could benefit from improved local specificity (Fig. 5a). To achieve this, we mutated and/or recombined regions of initial screening hits and used the fine-tuned Helix to predict their editing outcomes. We selected 12 with improved predicted bystander editing profiles for experimental testing (Fig. 5c).

The fine-tuned Helix accurately predicted the editing outcomes of the new gRNAs (R = 0.99, root mean squared error (RMSE) = 1.53), outperforming the non-fine-tuned Helix results for GRIK2 gRNAs (R = 0.88, RMSE = 5.36). An XGBoost model that is only trained using the fine-tuning data serves as another baseline (R = 0.90, RMSE = 4.18). Although all three models achieved relatively high Pearson correlations (R), the markedly lower RMSE of the fine-tuned Helix indicates much tighter agreement with experimental editing fractions across the full 0– 100% range. This accuracy is particularly important for bystander minimization, where small quantitative differences may carry meaningful functional consequences. Consistent with this, the fine-tuned Helix successfully predicted reduced bystander editing in the newly designed gRNAs, enabling further improvement of the lead candidate pool through iterative design.

## Discussion

In this study we highlight Helix, a task-specific transformer model that is highly predictive of ADAR-mediated RNA editing. Its improved accuracy arises from two attributes: a transformer architecture that efficiently leverages imbalanced training data and a modified attention mechanism that incorporates RNA secondary structure. By pairing Helix with DeepREAD, we created DeepHelix, an integrated NS distillation framework in which Helix scores large in silico gRNA design pools and the predicted editing outcomes are used to further train the DeepREAD diffusion models. As we will discuss, this combination allows for a flexible ADAR-recruiting gRNA design workflow.

Our primary aim was threefold: to identify a model architecture that can (i) embed structural knowledge of ADAR activity; (ii) scale with sparse and imbalanced data that is generated during screening; and (iii) benefit from improved performance after fine-tuning to enable iterative design during the discovery process. The practical limitations of LLMs and foundation models, such as extremely high data and compute needs, motivated us to evaluate whether a smaller, task-specific transformer could exhibit similar benefits while remaining practical to build and execute.

Although Helix is not a foundation model, given its narrow task, supervised training regime, and modest dataset, it demonstrates strong scaling behavior. Helix continues to improve when trained on single-target libraries containing tens of thousands of structurally diverse gRNAs, a setting in which XGBoost and convolutional neural network (CNN) baselines fail to benefit. While Helix performs comparably to these models when trained only on the well-balanced PolyTarget library, it uniquely leverages additional, highly imbalanced datasets, making it well-suited for iterative model updates during drug discovery. We speculate that both the use of the transformer architecture and the use of secondary structure attention enable the scaling of Helix. Although the single-target libraries are not diverse in sequence, they do possess significant structural diversity. Focusing attention maps based on structural predictions can allow the model to learn from the relationship between structure and editing despite the limited sequence diversity present in single-target libraries. These scaling properties pair well with advances in agentic AI and laboratory automation, which can continuously generate and integrate new empirical data^41,42^. Frameworks like DeepHelix have the potential to evolve rapidly and accelerate the prioritization of lead development candidates across multiple programs in parallel.

While scaling improves the global capabilities of Helix across targets, fine-tuning produces localized improvements with minimal adjustments to the pretrained backbone. Helix responds remarkably well to small amounts of labeled data: ∼100 in-cell observations for the GRIK2 TIS target were sufficient to fine-tune the model and achieve a predictive accuracy of R = 0.99, far exceeding its performance without fine-tuning. This is particularly important because Helix is trained on a biochemical assay that does not capture the full complexity of the cellular environment and the full gRNA-target duplex formed in cells. As previously mentioned, the model generates 45 base pair (bp) sequence, requiring us to extend the gRNA on both the 5’ and 3’ end to achieve the > 80 nucleotides required to recruit endogenous ADAR for DNA encoded gRNAs^8,9^. Additionally, the editing outcome in the cellular environment depends on ADAR isoform abundance, competition with other RNA-binding proteins, gRNA expression and processing, and the secondary structure of the endogenous pre-mRNA target region^5^. Fine-tuning Helix with data from tissue-relevant model systems enables the impact of the native editing environment and the full gRNA-target duplex to be incorporated into the model, improving its relevance for therapeutic design.

Because Helix is trained purely as a predictive model, it does not generate new sequences. This capability is provided instead by DeepREAD. Most LLMs derive their generative capacity from objectives such as autoregressive next-token prediction or masked language modeling during pre-training^43,44^, which capture statistical sequence patterns but are not designed to learn functional determinants, such as ADAR editing. In contrast, Helix uses attention to learn positional and structural dependencies across the gRNA–target duplex, enabling accurate per-adenosine predictions but no capacity for sequence exploration. To enable function-guided generation, we therefore adopt a NS distillation framework in which Helix provides functional pseudo-labels and DeepREAD, a diffusion model that learns the distribution of viable gRNA designs, generates new sequences aligned with Helix’s predictive landscape.

This framework preserves the strengths of both model types. Helix makes structure-aware ADAR editing predictions, while diffusion sampling maintains sequence diversity and allows explicit control over the structural features introduced during in silico sample preparation, such as maximizing cross-allelic activity or suppressing bystander editing at sensitive positions. Diffusion models retain these structural priors during sampling, whereas Helix identifies and corrects problematic sites through per-adenosine scoring. Generated sequences are subsequently rescored and passed through a flexible selection step incorporating additional criteria such as cross-reactivity, MFE, and sequence constraints (e.g., avoiding restriction enzyme sites or CpG motifs). Because generation, inference, and sampling are computationally lightweight, this pipeline supports rapid, iterative exploration of design space and yields a high-quality pool of candidates tailored to target-specific needs.

Training Helix from scratch with full control over the architecture offers the unique advantage of tightly coupling structural priors with learned representations, enabling Helix’s embeddings to encode features such as nucleotide pairing context and editing propensities essential for modeling ADAR activity. In Helix, this is achieved by incorporating the BPM directly into the attention mechanism, which causes the embeddings to encode structural features of the gRNA–target duplex. Looking ahead, embedding-based representations provide a promising avenue for further expanding model capability. Diffusion models can operate directly in embedding space^45^ – a strategy shown to improve sample diversity in protein design – and future iterations of DeepHelix could leverage this to generate gRNAs conditioned on structural or biochemical constraints learned by Helix, including those relevant for chemically modified gRNAs or antisense oligonucleotides. Embeddings that integrate additional contextual information (e.g., intramolecular gRNA folding, endogenous pre-mRNA structure, RNA-binding protein interactions, or chemical modifications) may yield richer representations for both prediction and generation, enhancing the biological knowledge and translational applicability of the model.

Many factors beyond intrinsic editing efficiency (e.g., gRNA expression, processing, global specificity, and safety) ultimately determine the clinical viability of an RNA editing therapeutic. The current workflow enables rapid identification of promising gRNAs in disease-relevant systems, with fine-tuning providing a mechanism to further optimize efficiency and specificity. Continued improvements in screening strategies and model design could further increase the likelihood that zero-shot predictions yield therapeutically viable gRNAs. For example, in silico prediction of off-target hybridization sites throughout the transcriptome, paired with Helix scoring, could proactively identify designs at risk of off-target editing. On the experimental side, we have previously used split-pool single-cell barcoding to optimize gRNA expression cassettes^14^, and similar high-throughput approaches could be extended to assess gRNA performance in disease-relevant cell types or even in vivo as genome-integration and single-cell sequencing technologies mature^46,47^. Larger data sets from disease relevant systems would provide valuable feedback to DeepHelix. Future models may incorporate richer embedding representations that capture additional biological context, enabling more nuanced modeling of the full gRNA-target structure, target accessibility, and cellular determinants of RNA editing.

Meanwhile, regulatory guidance is evolving to drive development of drugs for rare and ultra-rare diseases. The FDA’s Plausible Mechanism Pathway may allow a single IND to be expanded to cover multiple antisense therapeutics that target different mutations within the same gene^48^. Under this framework, once the foundational safety and efficacy package is established for a lead gRNA, subsequent gRNAs targeting additional mutations may require only streamlined characterization focused on on-target activity and off-target liabilities in relevant cell-based systems. In this context, rapid in silico design and testing of small gRNA panels to select a candidate for clinical testing could greatly broaden the number of patients eligible for treatment. We envision a future in which advanced computational screening tools such as DeepHelix, combined with efficient experimental validation and increasing regulatory flexibility, converge to accelerate the development of personalized RNA-based therapeutics enabling patients with urgent unmet needs to benefit from gene-editing technologies much sooner without compromising safety or efficacy.

## Methods

### Model architectures and training

Helix is a transformer encoder-decoder based model (Fig. 1), with a 12-layer encoder block and 12-layer decoder block. Each layer contains 20 attention heads, with modified scaled dot-product attentions and 5,120 as the dimension for the position-wise feed-forward network (FFN).

The input to the model is a one-hot encoded RNA sequence with length *l* as input *X* ∈ {0,1}^*l*×4^. The context of the input sequence is constructed by concatenating the target sequence, a linker loop (AAAAA throughout this study), and a gRNA sequence. To handle data loading in batches, we always pad the target sequence to length *l*_*t*_ by prepending ‘N’ bases at the beginning of the target, and then pad the concatenated sequence to length *l* by appending ‘N’ bases at the end of the sequence.

After the input sequence is converted into one-hot encoding, the model first maps it into embedding space with dimension *d* = 640 from the 4-dimensional one-hot encoded matrix through a learnable linear transformation:

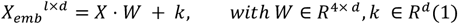

Then the model pushes *X*_*emb*_ through the Encoder network after adding a positional encoding and obtains an intermediate embedding of the entire sequence *X*_*intermediate*_ ∈ *R* ^*l* ×*d*^. To incorporate the secondary structural information into the modeling, we modified the scaled dot-product attentions in the Encoder network by making an addition of a Base-pairing Probability Matrix (BPM) to the attention map. Here we define the BPM as a symmetric matrix *B* ∈ [0,1]^*l*×*l*^ with each entry *b*_*j,i*_ = *b*_*i,j*_ ∈ [0,1] representing the pairing probability of the *i*^*th*^ and *j*^*th*^ base pairs of the sequence. The BPM is computed with EternaFold^33^ using default parameters, and added to the attention map of each attention head from all the layers within the Encoder network, resulting in a modified multi-head attention:

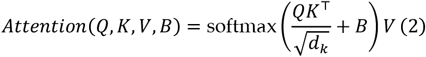

For the calculations of *Q, K, V* and the rest parts of transformer encoder layers, we follow the same procedures as described in the original implementation^29^; specifically we use the *X*_*emb*_ for *Q, K, V* matrices and BPM for *B* passing the secondary structural information.

After getting the intermediate embedding from:

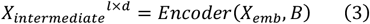

We then divide it into target and gRNA sequence embeddings by slicing the *X*_*intermediate*_ tensor at the token dimension at position *l*_*t*_, which results in a cutoff at the length of target sequence 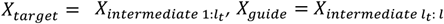. The target embedding *X*_*target*_ is then used as the query ( *Q* ) matrix, and the guide embedding *X*_*guide*_ is used as key (*K*) and value (*V*) matrices to the multi-head cross-attentions of the Decoder network, producing a final embedding *X*_*output*_ with the same length *l*_*t*_ as the target sequence. The Decoder network consists of 12 standard transformer decoder layers with vanilla cross-attentions, each with 20 attention heads, and using 5,120 as dimension for the position-wise FFN.

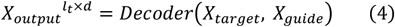

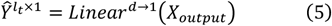

The output *X*_*output*_ from the Decoder network is then pushed through a final pointwise linear prediction head, to produce a scalar value of editing efficiency at each base pair *Ŷ*.

We use the standard MSE loss as objective function and standard gradient descent optimizer with momentum 0.9, learning rate 5e-5, and weight decay 1e-5 for 100 epochs. A linear learning rate scheduler with end factor 0.01 is used to adjust the learning rate during the training process. Our model is implemented with PyTorch.

### High-Throughput Sequencing datasets

The design, data generation, and preprocessing of the PolyTarget library, DeepREAD validation library, and single-target libraries are as previously described^17^. Briefly, all HTS libraries were in vitro transcribed, purified, and treated with recombinant ADAR in a biochemical reaction. Next, the libraries were subject to reverse transcription, library preparation, and next-generation sequencing (NGS) to quantify the percent of A-to-G conversion at each adenosine within the target sequence. For ease of data loading implementations, we imputed 0’s for all non-adenosine nucleotides within the target sequence, resulting in a numerical vector with values 0-100% and the same length for all target sequences.

### In-cell testing

For in-cell validation, DeepHelix generated gRNAs were extended to approximately 100 nucleotides by appending target-complementary antisense sequences to both the 5′ and 3′ ends of the gRNA sequence. Additional, internal loops are inserted near the extended junctions on both ends, which have been shown to enhance ADAR1 specificity^49^.

HEK293T cells were engineered to express the human GRIK2 minigene using the PiggyBac transposon system. For testing the mouse ortholog, the mouse Grik2 minigene plasmid was co-transfected with the plasmid encoded gRNAs. Cells were seeded 24 h before transfection in 96-well plates. Plasmids encoding the gRNAs of interest were transfected using TransIT-293 Transfection Reagent (Mirus #2704) according to the manufacturer’s instructions. Cells were harvested 48 h post-transfection, and RNA was isolated using the RNeasy Mini Kit (Qiagen #74104). Reverse transcription, PCR of the target region, and Sanger sequencing was performed with custom primers to quantify A-to-G conversion at each adenosine with the target region.

For comparison with the Helix predictions, we focused on the percent editing at target nucleotide positions present in the shorter target hybridization region of the gRNA prior to extension to ∼100 nucleotides.

SH-SY5Y cells were plated into 24-well plates at a density of 3.5e4 cells per well. The day after plating the cells were transduced with AAV encoding GRIK2 gRNAs at a ratio of 5E3 vg (viral genomes) per cell and RNA was harvested 7 days post transduction. Human iPSCs containing a doxycline inducible NeuroD1 expression cassette were differentiated into glutamatergic neurons over a period 10 – 14-days. After differentiation, cells were transduced with AAV encoding GRIK2 gRNAs at a ratio of 5e4 vg/cell and RNA was harvested 12 days post transduction. For both cell-types RNA was extracted with the MagMAX mirVana kit (Thermo # A25598). Reverse transcription, PCR of the target region, and Sanger sequencing was performed with custom primers to quantify A-to-G conversion at each adenosine within the target region.

### Evaluation and metrics

For benchmarking the zero-shot prediction performance of the models, we evaluated the performance on a withheld test dataset consisting of gRNAs from 21 targets unseen during the training phase (Fig. 2a). We filtered for structural integrity (i.e., proper self-annealing) and read depth > 50. After filtering, we subsampled 480 gRNAs for each target. For targets with less than 480 gRNAs after filtering, we included gRNAs that did not pass the filter to reach 480 samples.

We used Pearson and Spearman correlations as evaluation metrics for prediction performance. We computed the correlations between model predictions and observed on-target editing, as well as the observed editing at all adenosines present in the target sequence (both on-target and bystander adenosines).

For evaluation of the scaling law, we utilized the PolyTarget HTS library. At each given number of training targets, we used the data from the remaining targets for testing. Train/test targets were randomly shuffled 30 times to obtain mean and standard deviations of predictive performance. The model used is a XGBoost model (implemented with xgboost 1.7.4), fitted by grid search with cross validations.

### Noisy student training

Helix and DeepREAD were combined to create DeepHelix, a NS training workflow to generate and prioritize gRNAs to screen and validate in cells for therapeutic potential (Fig. 4a). For a given target of interest, we started by generating random designs (∼30,000 with possible duplicates). Random gRNAs were generated with a custom tool (GuideBuilder). We start with a canonical design (reverse complement of the target with an A-C mismatch feature at the on-target position). For each design, we randomly injected 0-4 features. The size (length of mismatch) of each feature is randomly sampled between 1-3, symmetrically or asymmetrically. GuideBuilder then leverages the ViennaRNA inverse fold function to identify a mutated gRNA sequence that produces the desired features within the gRNA-target duplex. We then use Helix to predict the editing outcome of the random gRNA designs. For each gRNA prediction, we computed the on-target editing efficiency and specificity (log ratio of on-target over total bystander edits) and used it as pseudo-labels for training the DeepREAD bit-diffusion model, a process similar to NS distillation^20,30,50^. DeepREAD is a conditional generative model – in each reverse diffusion process, it takes a target sequence, a set of desired editing profiles (on-target editing efficiency and specificity) as input and generates a gRNA sequence that is likely to lead to the desired editing profile. The NS training follows the same training procedure as described in original DeepREAD paper, except here we used synthetic data instead of experimental measurements – specifically, sequences are from random designs described above and the conditional editing profiles are from pseudo-labels generated by Helix combined with 10% relative random Gaussian noise. The noise level is determined by the empirical accuracy of Helix model from previous evaluations.

We used GRIK2 TIS as a target of interest to evaluate the performance of the DeepHelix. We synthesized random designs and generated pseudo-labels with Helix for training DeepREAD. After training the NS bit-diffusion generative model, we positively sampled the model for novel gRNA designs by conditioning on the GRIK2 TIS target sequence and editing values randomly sampled between 70-110% and specificity between 0.7-1.1. We then scored these generated designs with Helix, along with the random samples generated previously for training the NS bit-diffusion model, and prioritized these designs based on on-target editing and max bystander editing efficiencies.

### Evaluation of Embeddings

The embeddings used for evaluations are the intermediate outputs from the encoder layer of the model (Methods, equation (3)). For evaluating sequence embeddings from Helix, we generated a dataset containing 5,000 therapeutically related targets with 600 random gRNA designs for each target using GuideBuilder following the method described above. For visualizing the embedding landscape of gRNAs from different targets, we first conducted mean pooling along the sequence dimension and then apply non-linear dimension reduction methods like t-distributed Stochastic Neighbor Embedding (t-SNE) or UMAP to reduce the embedding dimension from *d* = 640 to 2. For better interpretations, we also performed linear projections with principal component analysis (PCA) on the embeddings after mean pooling along the sequence dimension, and computed Pearson correlations between top PCs and predicted properties of the sequences, such as MFE, on-target editing efficiencies, and max bystander editing efficiencies.

### Visualization of Attention Maps

To visualize the attention maps of Helix, we generated heatmaps for the attention weights from the last layer of the transformer encoder. The heatmap is generated by gathering the outputs across all attention heads from the last self-attention layer and computing the average pointwise attention score at each position across all attention heads.

### Designing gRNAs with cross-species reactivity

To design gRNAs with high cross-species editing efficiency and specificity, we developed a modified DeepHelix workflow. We generated a novel seed sequence for injection of random features with GuideBuilder by selectively using wobble base pairs to enable cross-species hybridization at divergent positions where it was possible to form wobble or canonical basepairs between species. Helix was used to generate pseudo-labels for editing outcomes for both species to train DeepREAD. As we didn’t augment the model architecture, it still takes one target sequence as conditional inputs. In this situation, we used the reverse complement of the pseudo-canonical gRNA as the target sequence for the NS bit-diffusion model. After model training, we positively sampled gRNA designs with high on-target editing and specificity for both species. These generative gRNA designs are combined with random samples, paired with both target sequences and scored by Helix to select designs with high predicted on-target editing and low maximum bystander editing for both species.

### Fine-tuning Helix with data from in-cell experiments

To utilize the in-cell experimental results and fine-tune the Helix model, we processed a batch of 62 designs tested in HEK293T cells into the same structure as the HTS data and used it to fine-tune a Helix model for predicting in-cell editing outcomes for GRIK2 TIS target. We loaded the model weights from a pretrained Helix model (for ADAR1), and during fine-tuning we used learning rate 5e-5 to train the model for 1000 epochs. The rest of the hyperparameters remain the same as the training phase. In this case, gRNAs were not generated by GuideBuilder, but were generated by introducing mutations or recombining top performing designs from the first batch. These designs were then scored using the fine-tuned model and designs with predicted high on-target editing and low max bystander editing were prioritized for in-cell validation. The fine-tuned model was then used to score and prioritize another batch of gRNA designs for the same target. For benchmarking, we scored the same gRNA designs with the non-fine-tuned Helix and an XGBoost model trained with the same 62 gRNA designs. We reported the RMSE and Pearson correlation for all adenosines (including on-target and all bystanders) for all three methods.

## Acknowledgements

We thank the following individuals for their support and contributions to this work: Adrian Briggs for scientific leadership and review of the manuscript; Ashley Benson, William Johnson, Dee Patel, and Emma Thuline for viral production and QC; J Moon, Rachel Feiring, and Jana Henry for technical advice and support.

## Author Contributions

Yingxin Cao conceptualized and developed Helix and DeepHelix with oversight from Yue Jiang and Ron Hause; Lina Bagepalli led gRNA screening efforts and collaboration with the computational team with oversight from Brian Booth and Yiannis Savva; Sierra Collins, Kris Adams, Alex Letizia, Samantha Edwards, Lina Bagepalli, and Lucia Shumaker contributed to gRNA screening efforts; Joanne Boysen contributed to data analysis and filtering of top candidates; Aydin Abiar contributed to the codebase for DeepREAD and DeepHelix; Stephen Burleigh engineered GRIK2 cell lines; Yiannis Savva provided scientific guidance and led gRNA engineering. Yingxin Cao, Yue Jiang, Yiannis Savva, and Brian Booth wrote the manuscript. All authors have approved the manuscript.

## Competing Interests

All authors are current or former employees of Shape Therapeutics Inc. and YC, LRB, YAS, AA, RJH, BJB, and YJ are inventors on patents and/or patent applications based on the site-directed RNA editing methods and/or the machine learning models described in this work.The authors declare no other competing interests.

