## Supplemental Figures for "Helix: a structure-aware deep learning model for accurate prediction of A-to-I RNA editing by endogenous ADARs"

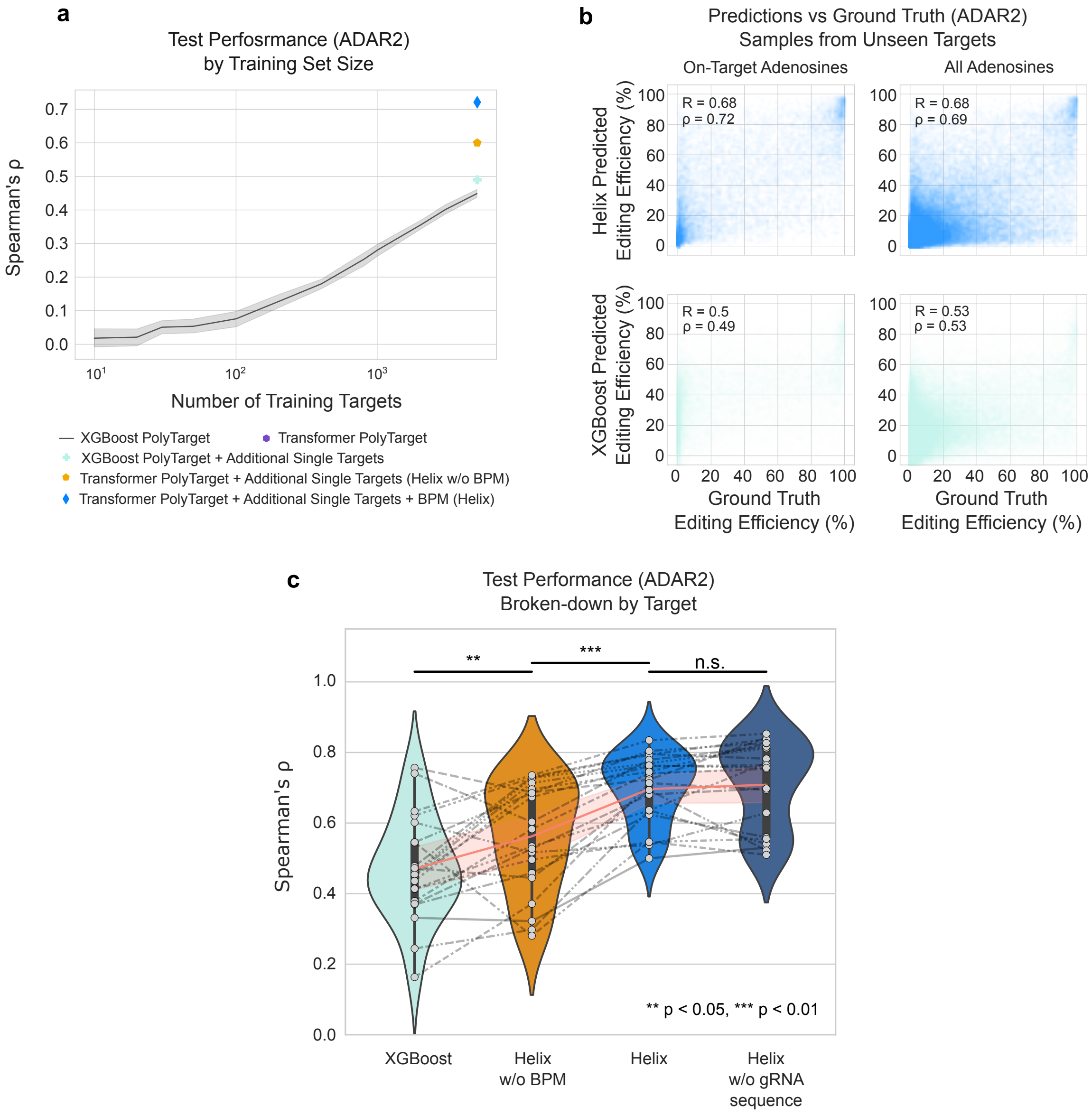


**Supplementary Fig. 1 | Model performance for ADAR2 editing outcomes. a,** Spearman correlation between test predictions and ground truth for ADAR2 editing outcomes, as a function of the number of targets used for training. **b,** Predictive performances on ADAR2 editing by Helix and XGBoost for on-target positions and all positions (bystander and on-target). **c,** Spearman correlation between predictions and ground truth for ADAR2 editing at on-target positions averaged within each target across different models.

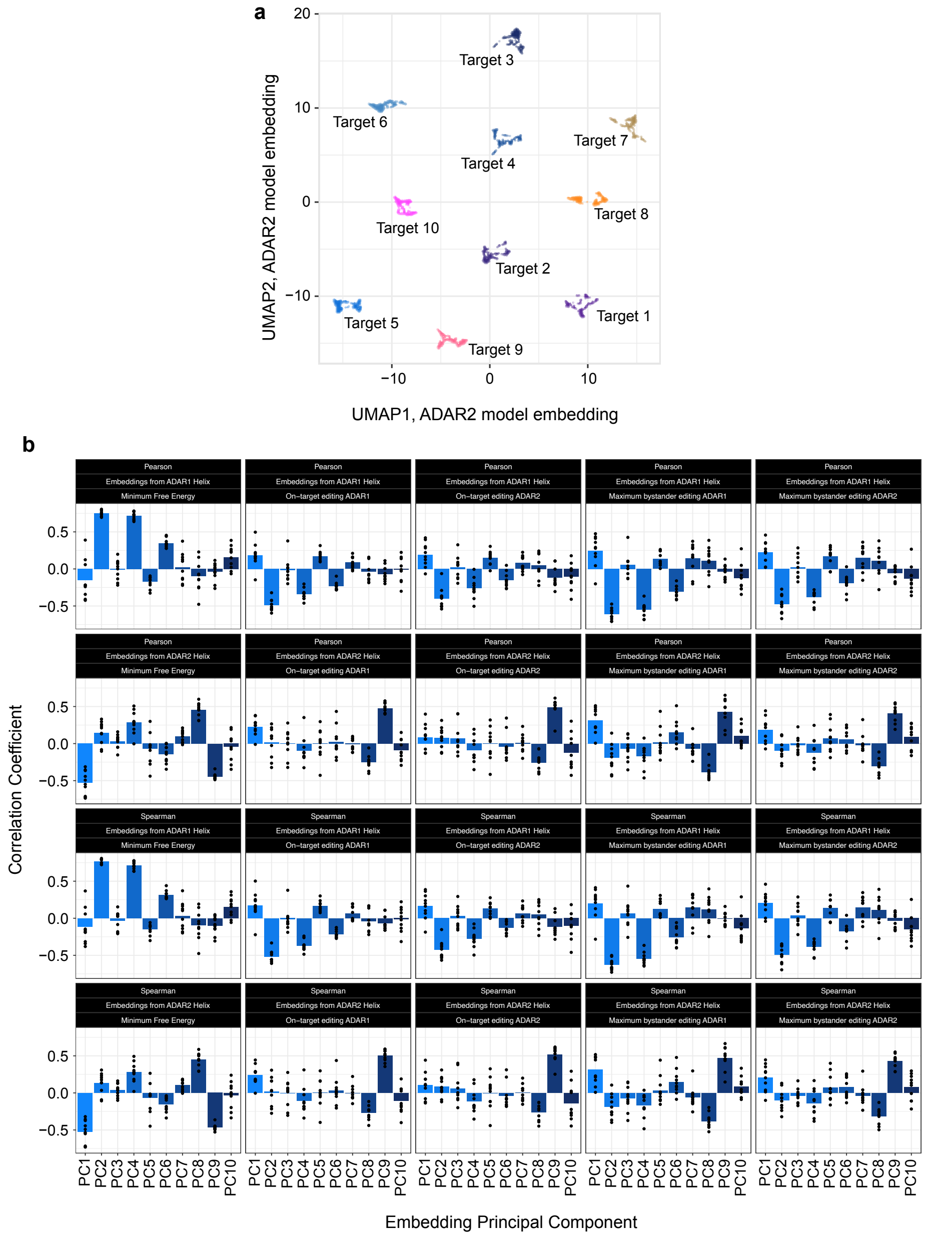


**Supplementary Fig. 2 | Model performance for ADAR2 editing outcomes. a,** 2-dimensional UMAP visualization of embeddings from the transformer encoder block of Helix ADAR2 model, for 10 targets with 600 gRNAs each. Each dot represents a gRNA sample. **b,** Pearson and Spearman correlation coefficients of the top 10 principal components (PC) of ADAR1 and ADAR2 embeddings with minimum free energy, on-target editing and maximum bystander editing.

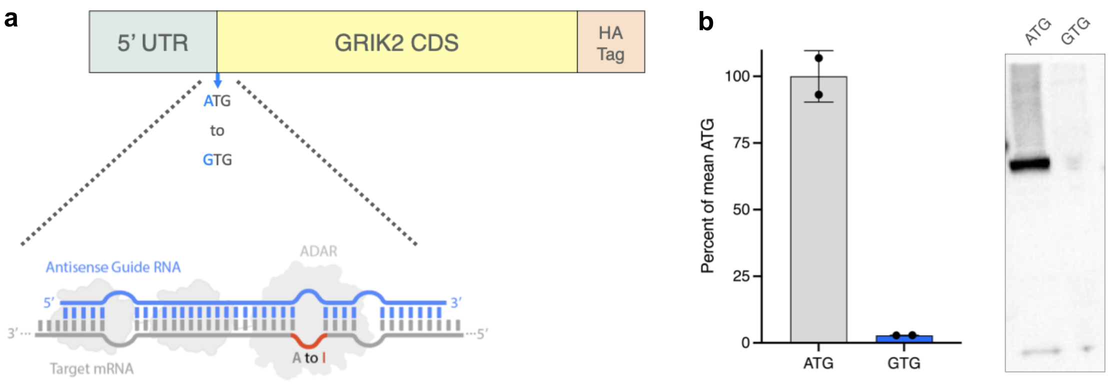


**Supplementary Fig. 3 l GRIK2 TIS target assessment. a,** Schematic of a mini-gene construct for validation that the conversion of an ATG to GTG at the GRIK2 translation initiation site (TIS) will abolish protein translation. The mini-gene contains a C-terminal HA-tag for ease of protein detection and is driven by an EF1a promoter with an SV40 polyA signal downstream. Also depicted is the site of gRNA hybridization where ADAR can be recruited to convert the TIS from ATG to ITG to abolish translation. **b,** To assess the impact of an A to G edit at the canonical TIS, mini-gene plasmids were generated with either an ATG or GTG as the start codon. These plasmids were then transfected into HEK293 cells and cell lysis was performed after 48 hours. Western blot was performed with 40 µg of protein input and exogenous GRIK2 was detected with an HA-tag primary rabbit mAb (cell Signaling cat# C29F4). Densitometry results from two replicates show the ATG-to-GTG mutation produced ~2.7% of the WT ATG signal.

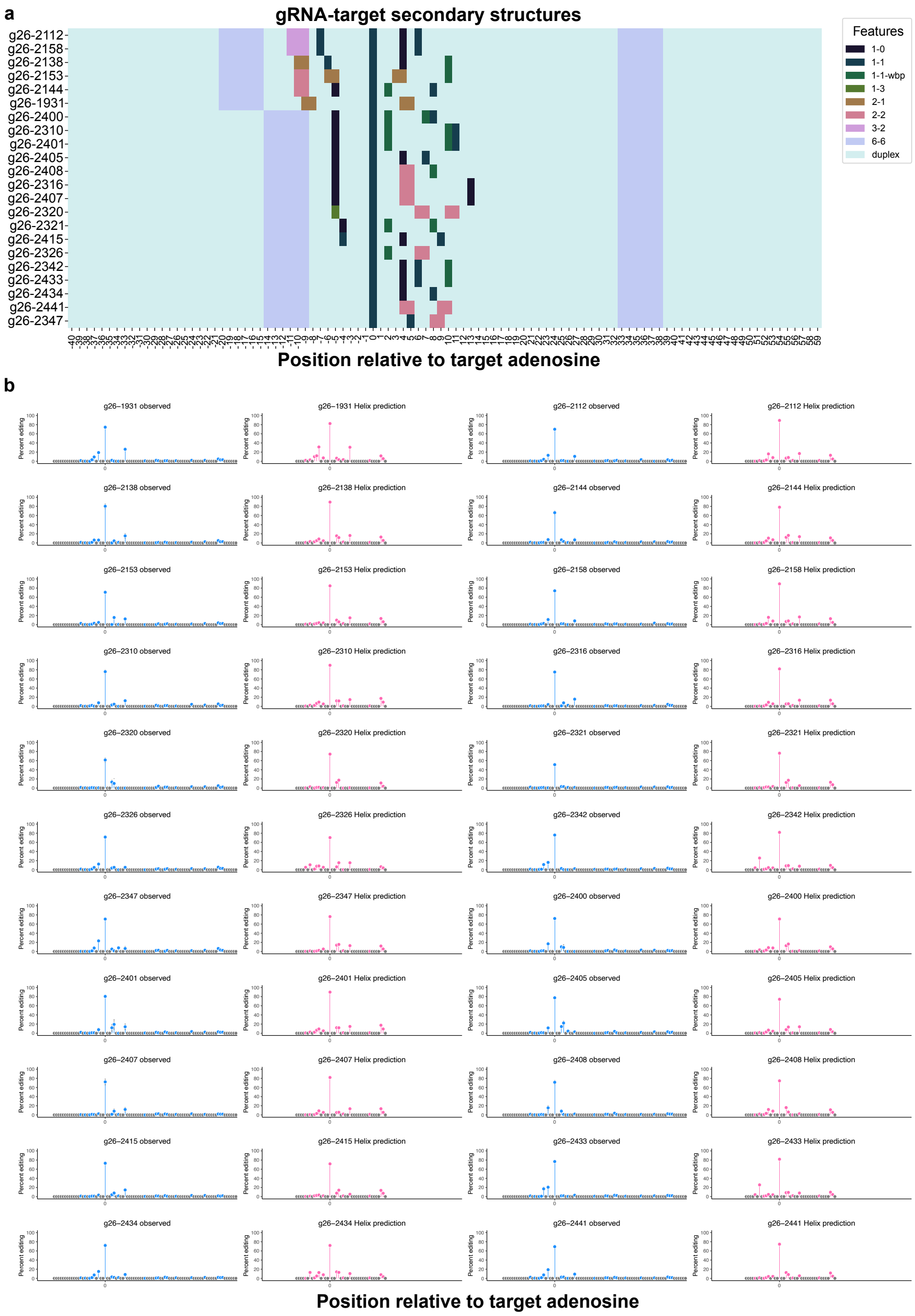


**Supplementary Fig. 4 | In-cell validation of Helix designs for GRIK2 TIS. a,** secondary structures of target-gRNA duplexes of top gRNAs validated in cells. **b,** Editing profiles observed in cells and predicted by Helix of top gRNA candidates.

**a**

```
Score:113 bits(125), Expect:8e−31,
Identities:85/100(85%),  Gaps:0/100(0%), Strand: Plus/Plus

Human GAAGTGGGTGCCGTGCGTGTGGGCACAGAAACACCATGAAGATTATTTTCCCGATTCTAAGTAATCCAGTCTTCAGGCGCACCGTTAAACTCCTGCTCTG
      ||||||||||||| ||| |  ||||| |||||| |||||||||||||| |||  || ||||||||| ||||||||| ||| || ||||| ||||||||||
Mouse GAAGTGGGTGCCGCGCGCGCAGGCACGGAAACATCATGAAGATTATTTCCCCAGTTTTAAGTAATCTAGTCTTCAGTCGCTCCATTAAAGTCCTGCTCTG
```

**b**

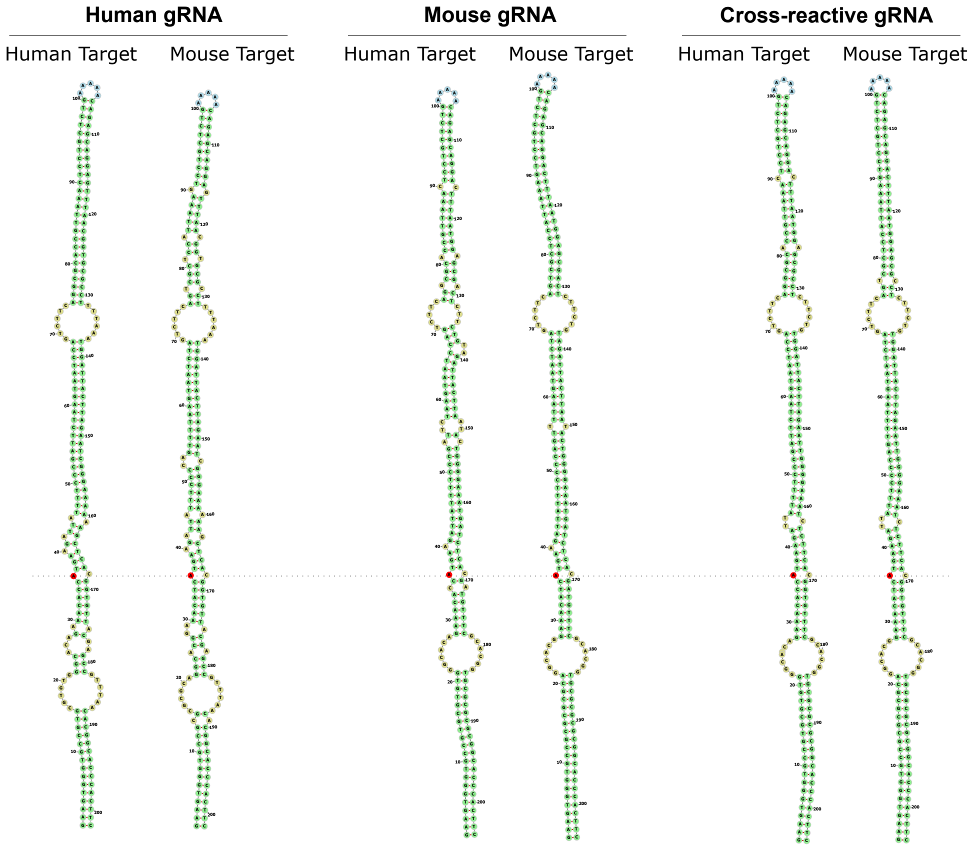


**Supplementary Fig. 5 | Cross-species reactive gRNA design. a,** alignment of GRIK2 TIS target sequences from human and mouse. **b,** secondary structures of a top human gRNA design, a top mouse gRNA design and a top cross-reactive gRNA design, in complex with human and mouse targets.
